# Single-particle tracking reveals neuronal activity-dependent shuttling of ARC/ARG3.1 protein between cytoplasmic clusters and the nucleus

**DOI:** 10.64898/2026.08.18.745435

**Authors:** Anna Dommarsnes Abrahamsen, Hauk Fevang, Yongyue Qian, Valentina Gandin, Zhe J. Liu, Ilaria Testa, Clive R. Bramham

## Abstract

The activity-regulated cytoskeleton-associated protein (ARC/ARG3.1) is a key regulator of synaptic plasticity and has both synaptic and nuclear functions. ARC is known to undergo nuclear import and export, yet the dynamic transport behavior of individual ARC particles remains unknown. Using live-cell single-particle tracking, we directly visualize ARC nucleocytoplasmic transport and shuttling in primary hippocampal neurons. Synaptic activation by chemical long-term potentiation (cLTP) treatment increases shuttling behavior and reveals a previously underappreciated organization of ARC within the neuronal cell body cytoplasm, characterized by perinuclear ARC clusters. Disruption of the N-terminal ARC oligomerization motif markedly reduced both perinuclear cluster formation and nucleocytoplasmic shuttling. Together, these findings reveal an activity-dependent relationship between ARC self-assembly, perinuclear organization, and nucleocytoplasmic trafficking, providing a potential mechanism for coordinating the synaptic and nuclear functions of ARC during neuronal plasticity.

## Introduction

Long-lasting forms of synaptic plasticity require activity-dependent gene transcription, necessitating communication between activated synapses and the nucleus to drive persistent changes in neuronal function(1) (1–3). Immediate-early genes (IEGs) are central mediators of this synapse-to-nucleus signaling, rapidly translating neuronal activity into molecular programs that support learning and memory. Among these, activity-regulated cytoskeleton-associated protein (ARC, also known as ARG3.1) has emerged as a key regulator of synaptic plasticity and memory consolidation (4–7). ARC is predominantly expressed in glutamatergic neurons, where it can be detected in the nucleus and throughout the somato-dendritic compartment of activated neurons. At synapses, ARC regulates AMPA receptor trafficking and actin remodeling to regulate both long-term depression (LTD) (8–11) and long-term potentiation (LTP) (12, 13). In the nucleus, ARC has been implicated in regulation of chromatin state and nuclear speckle formation (14–19). Despite these diverse roles, most studies have examined ARC’s synaptic and nuclear functions in isolation, leaving unresolved whether and how ARC activity is coordinated across subcellular compartments.

The ability of ARC to function both at synapses and within the nucleus raises a fundamental question of how these spatially separated activities are coordinated. Communication between the cytoplasm and nucleus is mediated through the nuclear pore complex, which regulates molecular exchange across the nuclear envelope. Although small proteins may diffuse passively through nuclear pores, regulated communication between the cytoplasm and nucleus typically relies on proteins that undergo signal-dependent nucleocytoplasmic transport. Such shuttling proteins dynamically redistribute between compartments in response to extracellular cues, thereby coupling cytoplasmic signaling events to changes in nuclear gene expression. Defects in nucleocytoplasmic transport have been implicated in numerous neurological and neurodegenerative disorders, highlighting the importance of this process for neuronal function (20, 21).

ARC possesses a functional nuclear localization signal, a nuclear retention domain, and a nuclear export signal, and previous studies have demonstrated that neuronal activity regulates its nucleocytoplasmic localization (15, 22). While demonstrating nucleocytoplasmic transport, existing evidence is derived primarily from measurements of steady-state protein localization in fixed cells or population-level imaging approaches. Consequently, it remains unknown whether individual ARC particles dynamically shuttle between the cytoplasm and nucleus in living neurons. Resolving this question is essential for understanding how ARC regulates nuclear responses during activity-dependent plasticity.

Here, we combined live-cell single-particle tracking microscopy with quantitative image analysis to directly visualize the movement of individual ARC particles in primary hippocampal neurons. We identify ARC as a bona fide nucleocytoplasmic shuttling protein whose transport is increased following chemically induced long-term potentiation (cLTP). Furthermore, we show that neuronal activity promotes the formation of perinuclear ARC clusters that are preferentially associated with shuttling trajectories. Finally, we demonstrate that disruption of ARC oligomerization markedly reduces both perinuclear cluster formation and nucleocytoplasmic shuttling, revealing a previously unrecognized role for ARC self-assembly in regulating activity-dependent nucleocytoplasmic transport.

## Results

### ARC nuclear crossings are increased following neuronal stimulation

To examine ARC nucleocytoplasmic transport under steady-state conditions, primary hippocampal neurons expressing Halo-tagged ARC from a constitutive CMV promoter were imaged using single-particle tracking (SPT, Fig. 1A). Sparse labeling with a low concentration of Janelia Fluor Halo ligand generated spatially isolated fluorescent ARC particles that could be unambiguously localized and linked into trajectories over time. SPT is a technique that enables real-time monitoring of individual particles in living cells and is well suited for studying molecular transport across the nuclear envelope (23, 24). To identify nucleocytoplasmic transport events, we developed a stringent core-crossing analysis pipeline that excludes apparent crossing events arising from transient interactions with the nuclear border that do not represent true compartment transitions (Fig. 1B).

**Figure 1:**
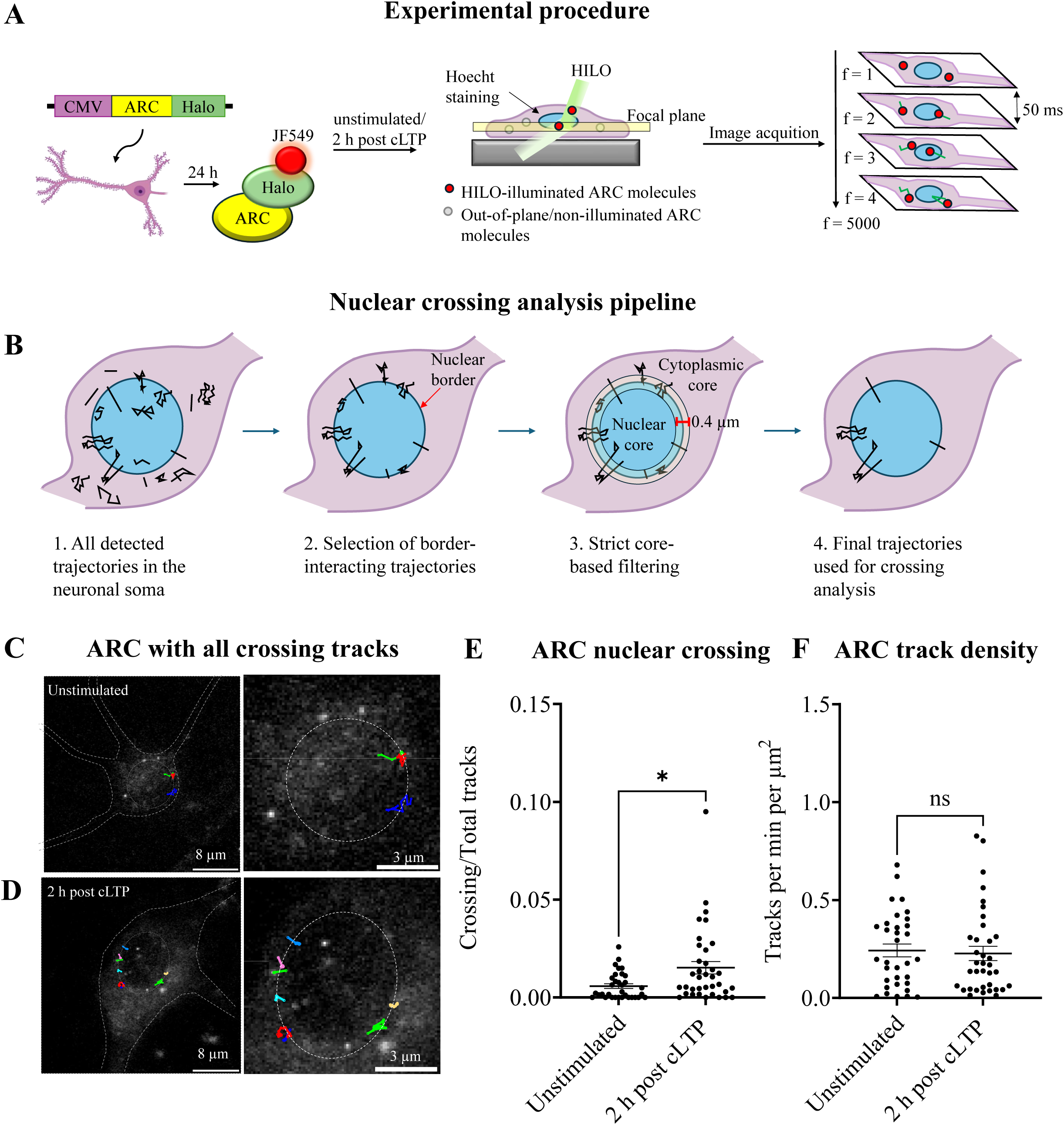
ARC nuclear crossings increase following cLTP treatment. **(A)** Experimental workflow for single-particle tracking (SPT) of ARC–Halo. Primary hippocampal neurons (DIV14–18) expressing ARC fused to a C-terminal HaloTag (ARC–Halo) were imaged under unstimulated conditions or 2 h following chemical long-term potentiation (cLTP) induced by glycine, bicuculline, and strychnine. Individual ARC molecules were labelled with JF549 HaloTag ligand and visualized by HILO microscopy. The nucleus was identified using Hoechst staining. Movies consisted of 5,000 frames acquired at 50 ms intervals. **(B)** Schematic of the stringent core-compartment analysis used to identify nuclear crossing events. Border-interacting trajectories were first identified as candidate crossings. To minimize false-positive classifications arising from localization uncertainty or transient interactions with the nuclear envelope, nuclear and cytoplasmic core regions were defined as localizations occurring at least 0.2 µm from the nuclear border. Only trajectories exhibiting transitions between the cytoplasmic and nuclear core compartments were classified as strict nuclear crossing events. **(C-D)** Representative unstimulated neuron (top) and neuron 2 h following cLTP (bottom) showing trajectories classified as strict nuclear crossing events. One representative frame from each movie is shown (left; scale bars, 8 µm). Detected crossing trajectories are displayed in different colours. Right panels show enlarged views of the nucleus with overlaid crossing trajectories (scale bars, 3 µm). **(E)** Fraction of trajectories undergoing strict nuclear crossing per cell in unstimulated neurons (0.0058 ± 0.0011) and 2 h following cLTP (0.0157 ± 0.0032). cLTP significantly increased the fraction of trajectories undergoing strict nuclear crossing (Mann–Whitney test, *P* = 0.0129). Each point represents one cell; bars indicate mean ± s.e.m. Unstimulated: *n* = 33 cells; 2 h post-cLTP: *n* = 36 cells from six independent neuronal cultures. **(F)** ARC trajectory density (tracks min⁻¹ µm⁻²) per cell in unstimulated neurons and neurons 2 h following cLTP. No significant difference in trajectory density was observed between conditions (Mann–Whitney test, *P* = 0.5767), indicating that the increase in strict nuclear crossing events was not attributable to differences in the number of detected trajectories.

Using this approach, we detected nuclear crossing events of individual ARC particles in primary hippocampal neurons. Under unstimulated conditions, these events were infrequent, with most cells exhibiting few or no crossings. Given ARC’s roles in activity-dependent plasticity, we next asked whether neuronal stimulation alters nucleocytoplasmic transport. Neuronal cultures receiving chemical long-term potentiation (cLTP) treatment, using 10 min application of glycine, bicuculline, and strychnine (25), and imaged 2 h post-stimulation (Fig. 1D). Following cLTP treatment, the fraction of ARC trajectories undergoing nuclear crossing increased approximately threefold compared with unstimulated neurons (Mann–Whitney U test, *P* = 0.013; Fig. 1E). To determine whether this increase could simply reflect differences in the number of detected ARC trajectories, we quantified trajectory density on a per-cell basis. No significant difference in trajectory density was observed between unstimulated and stimulated neurons (Mann–Whitney U test, p = 0.58; Fig. 1G).

To determine whether the observed increase reflected nonspecific molecular diffusion or an analysis artefact, we performed identical experiments using an empty-Halo construct (Supplementary Fig. 1A–D). In contrast to ARC-Halo, cLTP did not alter the fraction of empty-Halo trajectories undergoing nuclear crossing (Mann–Whitney U test, p = 0.22, Supplementary Fig. 1C). Likewise, trajectory density remained unchanged between unstimulated and stimulated empty-Halo cells (Supplementary Fig. 1D). Together, these findings demonstrate that neuronal stimulation selectively increases nuclear envelope crossing of ARC rather than causing a global increase in molecular mobility or apparent crossing events. These results provide direct visualization of activity-dependent nuclear envelope crossing by individual ARC particles in living neurons.

### ARC is a nucleocytoplasmic shuttling protein

Having established that ARC undergoes nuclear crossing in living neurons, we next sought to determine the directionality of these transport events. Nuclear-crossing trajectories were classified into three categories: import, in which a particle moved from the cytoplasm into the nucleus; export, in which a particle moved from the nucleus into the cytoplasm; and shuttling, in which a particle underwent two or more nuclear crossings within a single trajectory (Fig. 2A). Although previous studies have shown that ARC is capable of both nuclear import and export (15), whether individual ARC particles repeatedly shuttle between the nucleus and cytoplasm has remained unknown.

**Figure 2.**
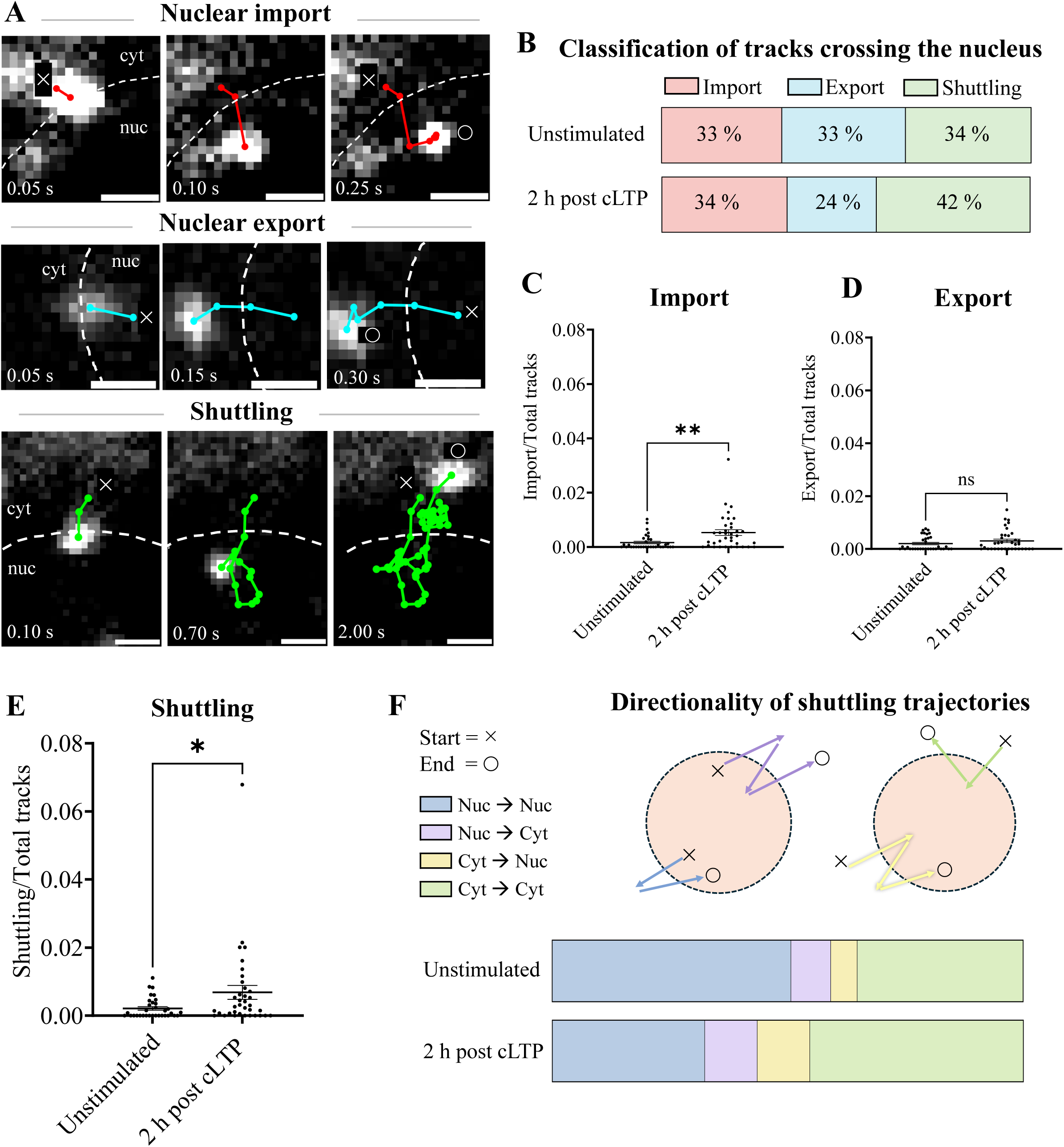
ARC undergoes bidirectional nucleocytoplasmic shuttling following synaptic stimulation. **(A)** Representative examples of individual ARC–Halo trajectories classified as nuclear import, nuclear export, or shuttling events. Import trajectories crossed from the cytoplasmic core to the nuclear core, whereas export trajectories crossed from the nuclear core to the cytoplasmic core. Shuttling trajectories underwent two or more strict core transitions between the nucleus and cytoplasm. Time stamps indicate elapsed time from the start of each trajectory. Dashed lines indicate the nuclear border. Scale bars, 0.8 µm. **(B)** Distribution of nuclear-crossing ARC trajectories classified as import, export, or shuttling events. Unstimulated, n = 209 trajectories; 2 h post cLTP, n = 471 trajectories. **(C-E)** Per-cell fraction of trajectories classified as import (C), export (D), or shuttling (E) in unstimulated neurons and 2 h following cLTP stimulation. The fraction of import trajectories increased following cLTP (*P* = 0.0012, Mann–Whitney U test; C), whereas the fraction of export trajectories was unchanged (*P* = 0.3827, Mann–Whitney U test; D). The fraction of shuttling trajectories also increased following stimulation (*P* = 0.0120, Mann–Whitney U test; E). Each data point represents one cell. Unstimulated, *n* = 33 cells; 2 h post-cLTP, *n* = 36 cells. **(F)** Classification of shuttling trajectories according to their start and end compartments in unstimulated neurons and 2 h following cLTP stimulation. Shuttling trajectories were subdivided into four categories according to the compartment in which they originated and terminated. In unstimulated neurons, 50% of shuttling trajectories were classified as nucleus-to-nucleus (Nuc→Nuc), 8% as nucleus-to-cytoplasm (Nuc→Cyt), 5% as cytoplasm-to-nucleus (Cyt→Nuc), and 35% as cytoplasm-to-cytoplasm (Cyt→Cyt). At 2 h post-cLTP, 32% were Nuc→Nuc, 11% Nuc→Cyt, 11% Cyt→Nuc, and 45% Cyt→Cyt.

Among all nuclear-crossing trajectories, import, export, and shuttling events occurred at comparable frequencies under unstimulated conditions (33%, 33%, and 34%, respectively; Fig. 2B). Following cLTP stimulation, the proportion of shuttling trajectories increased from 34% to 42%, whereas export trajectories decreased from 33% to 24%. The relative contribution of import trajectories among all nuclear-crossing events remained similar (34%; Fig. 2B). Next we quantified the fraction of each transport class on a per-cell basis. The fraction of import trajectories increased significantly following cLTP stimulation (Mann–Whitney U test, *P* = 0.0012; Fig. 2C), whereas export trajectories were unchanged (Mann–Whitney U test, *P* = 0.38; Fig. 2D). In addition, the fraction of shuttling trajectories increased significantly following stimulation (Mann–Whitney U test, *P* = 0.01; Fig. 2E), indicating that neuronal activity promotes repeated nucleocytoplasmic transport of individual ARC particles. We next quantified each transport class on a per-cell basis. Nuclear import events increased significantly following cLTP stimulation (Mann–Whitney U test, *p* < 0.01; Fig. 2C), whereas nuclear export events were unchanged (Fig. 2D). Notably, shuttling events also increased significantly following stimulation (Mann–Whitney U test, *p* < 0.01; Fig. 2E), demonstrating that neuronal activity promotes repeated nucleocytoplasmic transport of individual ARC particles.

Finally, to further characterize ARC shuttling behavior, we determined the compartment in which each shuttling trajectory originated and terminated (Fig. 2F). Under unstimulated conditions, half of all shuttling trajectories both originated and terminated within the nucleus, whereas 35% originated and terminated in the cytoplasm. Following cLTP stimulation, the proportion of Nuc→Nuc trajectories decreased from 50% to 32%, while Cyt→Cyt trajectories increased from 35% to 45%. Smaller increases were also observed for Nuc→Cyt trajectories (8% to 11%) and Cyt→Nuc trajectories (5% to 11%). Thus, neuronal stimulation shifted ARC shuttling from predominantly nucleus-confined trajectories toward trajectories that originated and terminated in the cytoplasm following transient nuclear entry. Together, these findings reveal that ARC undergoes dynamic bidirectional transport across the nuclear border and identify, to our knowledge, the first direct evidence that individual ARC particles actively shuttle between the nucleus and cytoplasm in living neurons.

### ARC forms clusters in the perinuclear region post neuronal stimulation

Because neuronal synaptic stimulation increased the nucleocytoplasmic transport of ARC, we next asked whether it also altered its intracellular distribution. ARC fluorescence intensity was quantified within the cytoplasmic, perinuclear, and nuclear compartments (Fig. 3A–C).

**Figure 3:**
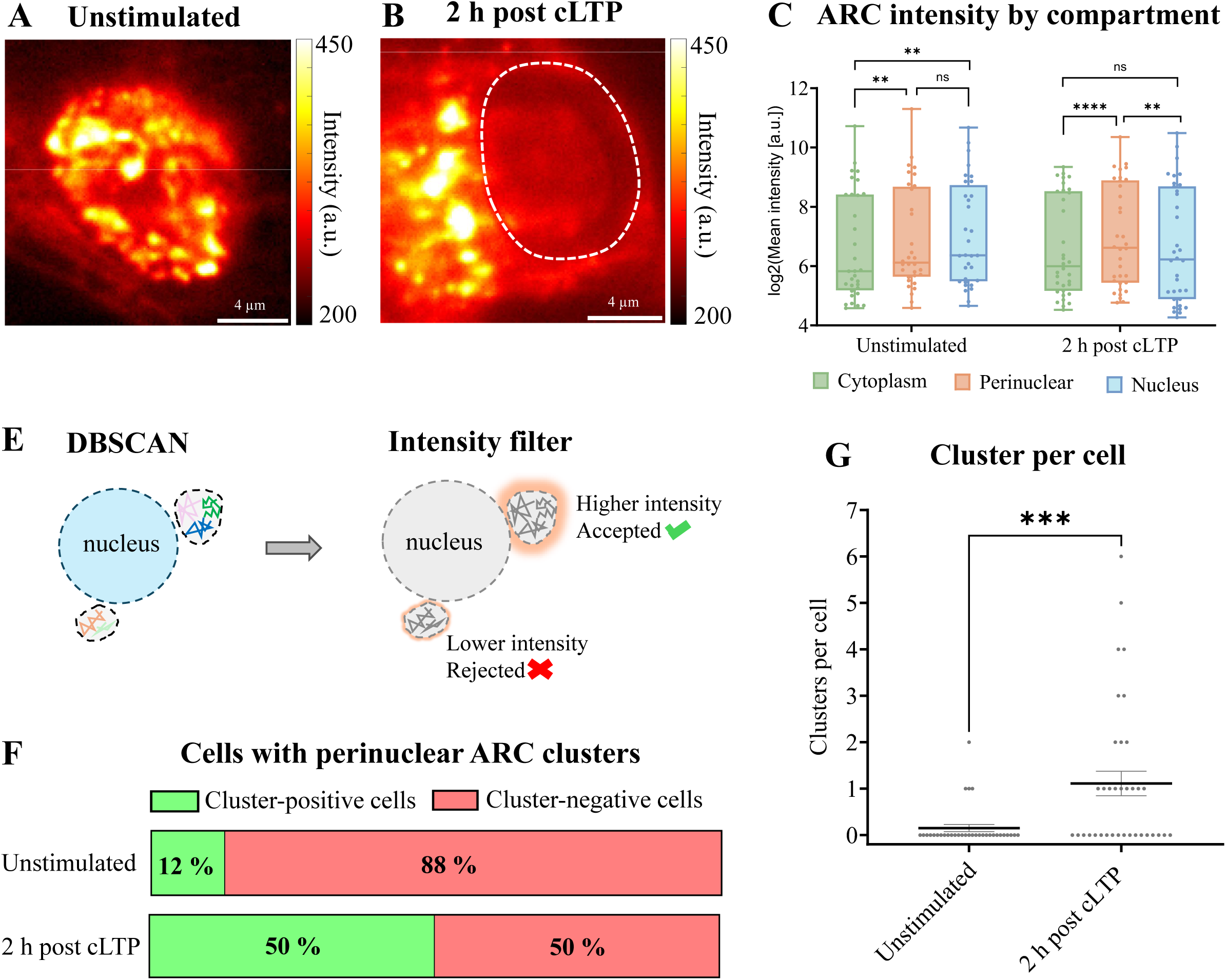
ARC forms clusters in the perinuclear cytoplasm after synaptic stimulation. **(A-B)** Representative maximum-intensity projections of ARC–Halo fluorescence in unstimulated neurons (A) and 2 h following cLTP stimulation (B). Dashed outline indicates the nucleus. Scale bars, 4 µm. **(C-D)** Quantification of ARC fluorescence intensity within the cytoplasm, perinuclear region, and nucleus. Under unstimulated conditions, ARC intensity was significantly higher in both the perinuclear region (*P* = 0.0022) and nucleus (*P* = 0.0022) than in the cytoplasm, whereas nuclear and perinuclear intensities were not significantly different (Friedman test with Dunn’s multiple-comparisons test). Following cLTP stimulation, ARC intensity was significantly higher in the perinuclear region than in both the cytoplasm (*P* < 0.0001) and nucleus (*P* = 0.0019), whereas nuclear and cytoplasmic intensities were not significantly different (P = 0.5846). Each data point represents one cell. Unstimulated, *n* = 33 cells; 2 h post-cLTP, *n* = 36 cells. **(E)** Schematic of the cluster detection pipeline. Candidate clusters identified by DBSCAN were subjected to an intensity-based validation step, and only clusters exceeding the predefined intensity threshold were retained for downstream analysis. **(F)** Percentage of cells containing at least one validated perinuclear ARC cluster. Unstimulated, 12% (4/33 cells); 2 h post-cLTP, 50% (18/36 cells). **(G)** Number of validated perinuclear ARC clusters per cell in unstimulated neurons and 2 h following cLTP stimulation. Cluster number increased significantly following stimulation (Mann–Whitney U test, *P* = 0.0011). Each data point represents one cell.

Under unstimulated conditions, ARC intensity was higher in both the perinuclear region and nucleus than in the cytoplasm, whereas perinuclear and nuclear intensities were comparable (Dunn’s multiple-comparisons test: cytoplasm versus perinuclear region, *P* = 0.002; cytoplasm versus nucleus, *P* = 0.002; perinuclear region versus nucleus, *P* > 0.99; Fig. 3C). Two hours following cLTP stimulation, ARC intensity was significantly higher in the perinuclear region than in either the cytoplasm or nucleus (cytoplasm versus perinuclear region, *P* < 0.0001; perinuclear region versus nucleus, *P* = 0.002), whereas cytoplasmic and nuclear intensities did not differ significantly (Fig. 3C). These findings indicate that neuronal stimulation redistributes ARC toward the perinuclear compartment.

Inspection of the fluorescence images further revealed discrete punctate ARC structures within the perinuclear region. To identify these structures systematically, we developed a cluster-detection pipeline combining density-based spatial clustering of applications with noise (DBSCAN) (26) with an intensity-based validation step that excluded low-intensity detections (Fig. 3E).

Using this approach, the proportion of cells containing at least one validated perinuclear ARC cluster increased from 13% under unstimulated conditions to 50% following cLTP stimulation (4/31 and 18/36 cells, respectively; Fig. 3F). Consistent with this increase, the number of validated perinuclear clusters per cell was significantly higher 2 h following cLTP induction (Mann–Whitney U test, *P* = Fig. 3G). No validated clusters were detected in empty-Halo control cells (Supplementary Fig. 2E-J), indicating that cluster detection required ARC–Halo signal. Together, these findings show that neuronal stimulation promotes the perinuclear enrichment of ARC and the formation of discrete perinuclear ARC clusters.

### Arc shuttling preferentially associates with perinuclear clusters

The emergence of perinuclear ARC clusters prompted us to investigate whether these structures are associated with nucleocytoplasmic shuttling. Because unstimulated neurons contained too few perinuclear clusters for reliable analysis, subsequent analyses were performed using neurons 2 h after cLTP stimulation. Representative time-lapse images illustrate a shuttling ARC particle that repeatedly associates with the same perinuclear cluster during nuclear entry and subsequent exit, exemplifying trajectories classified as cluster-associated shuttling (Fig. 4A). Mapping shuttling trajectories relative to perinuclear clusters further revealed that trajectories contacted some clusters, whereas others showed no detectable association (Fig. 4B).

**Figure 4.**
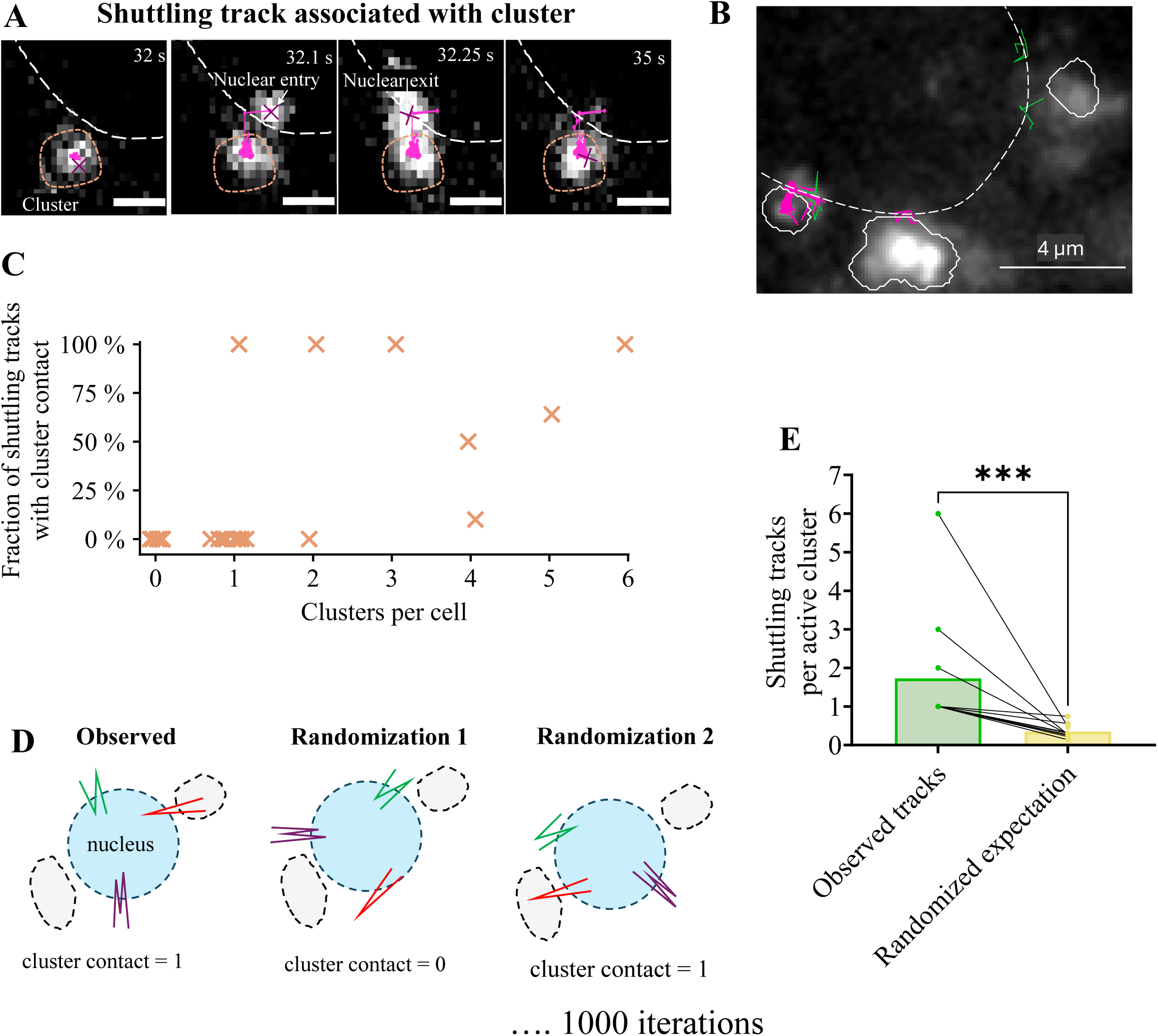
ARC shuttling events preferentially associated with perinuclear ARC clusters. **(A)** Representative single-molecule trajectory of a shuttling ARC molecule contacting a perinuclear ARC cluster. Consecutive frames show an ARC molecule (magenta) crossing the nuclear envelope and contacting a perinuclear ARC cluster (cyan arrows). Dashed outlines indicate the nuclear boundary (white) and the validated ARC cluster (red). Scale bars, 0.8 µm. **(B)** Representative spatial map of ARC shuttling trajectories and validated perinuclear ARC clusters in a neuron 2 h following cLTP stimulation. Shuttling trajectories are shown relative to the nuclear boundary (dashed white line) and validated perinuclear clusters (solid white outlines). Scale bar, 4 µm. **(C)** Relationship between the number of perinuclear ARC clusters and the fraction of shuttling trajectories associated with at least one cluster on a per-cell basis. Each point represents one cell. The fraction of cluster-associated shuttling events positively correlated with cluster abundance (Spearman correlation, r_s_ = 0.72, ***P = 0.0001, n = 23 cells). **(D)** Schematic of the randomization analysis. For each cell, the observed number of shuttling trajectories contacting a validated ARC cluster was compared with the mean obtained from 1,000 randomized realizations of the same trajectories. During each randomization, every shuttling trajectory was repositioned to a random location along the nuclear boundary while preserving its geometry and maintaining a valid shuttling event. Cluster contacts were rescored after each randomization, generating a null distribution against which the observed number of cluster-associated shuttling trajectories was compared. **(E)** Comparison of the observed fraction of shuttling trajectories contacting validated ARC clusters with the randomized expectation for the same cells. Each line connects the observed value and the mean of 1,000 randomized realizations from the same cell. Statistical significance was assessed using a paired Wilcoxon signed-rank test.

Across individual neurons, we calculated the fraction of total shuttling trajectories that contacted at least one perinuclear ARC cluster, thereby normalizing cluster-associated shuttling to the total number of shuttling events in each neuron. This fraction increased with the number of perinuclear clusters present in each cell (Spearman *r* = 0.72, *P* = 0.0001; Fig. 4C). Thus, neurons containing more perinuclear ARC clusters exhibited a greater proportion of their total shuttling trajectories in contact with clusters.

Because neurons containing more clusters also provide more opportunities for trajectories to encounter a cluster by spatial overlap alone, we next asked whether the observed trajectory–cluster contacts exceeded those expected by chance. To address this, shuttling trajectory positions were randomized and cluster contacts were quantified across 1,000 randomizations (Fig. 4D). We found that ARC shuttling was associated with a subset of perinuclear ARC clusters, with these clusters contacted by significantly more shuttling trajectories in the observed data than expected from randomized trajectory positions (Fig. 4E). Together, these findings indicate that ARC shuttling preferentially associates with discrete perinuclear ARC clusters and raise the possibility that this subset of clusters represents active sites involved in ARC nucleocytoplasmic trafficking.

### ARC oligomerization promotes perinuclear cluster formation and nucleocytoplasmic shuttling

ARC assembles into structures ranging from dimers to higher-order oligomers, including 32-mers and virus-like capsids (6, 27–34). To determine whether ARC oligomerization contributes to perinuclear cluster formation and nucleocytoplasmic shuttling, we compared ARC^WT^ with the oligomerization-deficient mutant ARC^s113–119A^ (27) in hippocampal neurons 2 h post stimulation. Representative images showed that ARC^WT^ formed distinct perinuclear clusters (Fig. 5A), whereas ARC^s113–119A^ displayed a more diffuse distribution with few detectable clusters (Fig. 5B). Consistent with these observations, the number of validated perinuclear clusters per cell was significantly reduced in ARC^s113–119A^-expressing neurons compared with ARC^WT^ (Fig. 5C), indicating that ARC oligomerization promotes efficient perinuclear cluster formation.

**Figure 5.**
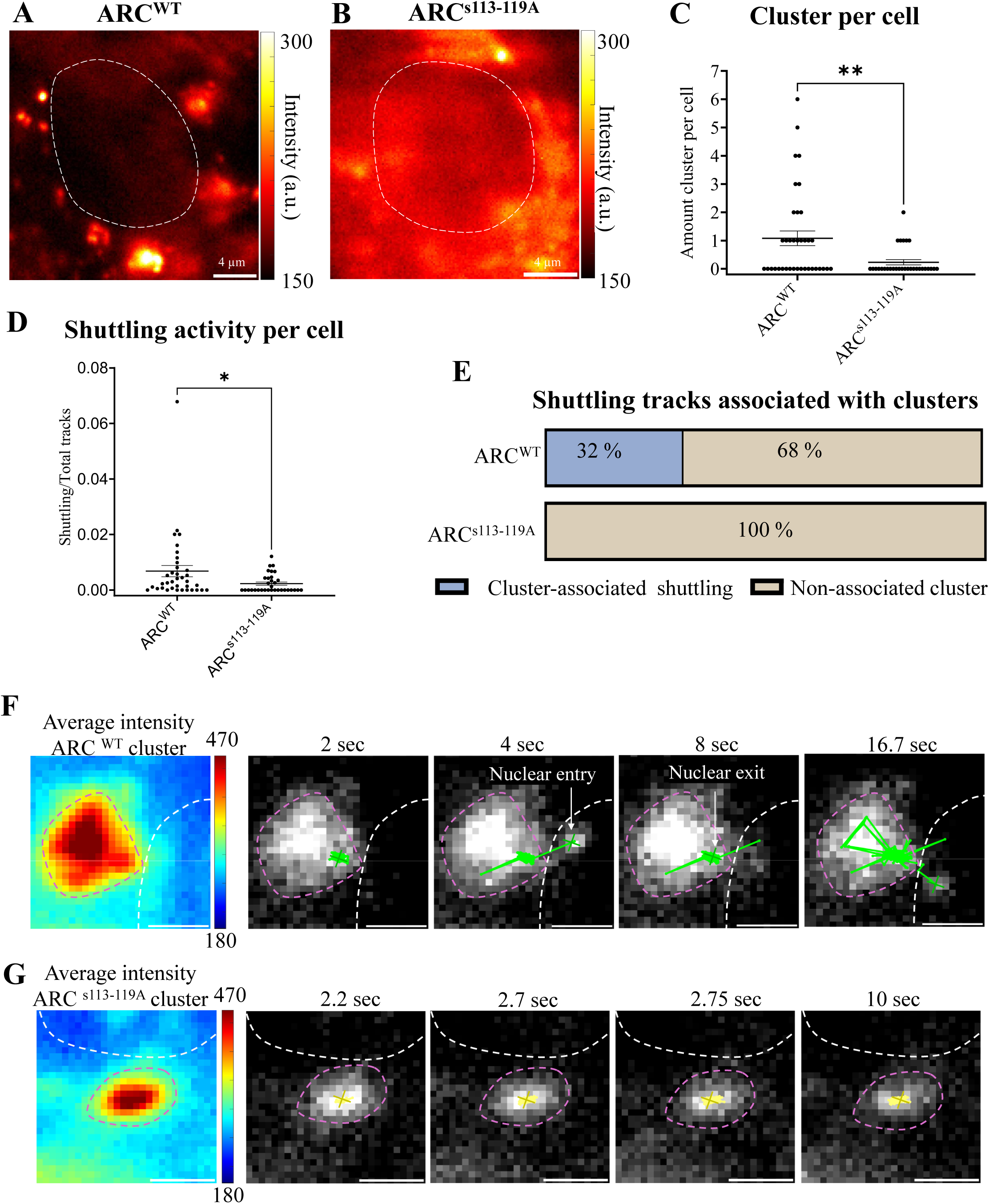
ARC oligomerization promotes perinuclear cluster formation and cluster-associated nucleocytoplasmic shuttling. **(A–B)** Representative average-intensity maps of ARC^WT^ (A) and the oligomerization-deficient mutant ARC^S113–119A^ (B). Dashed white lines indicate the nuclear boundary. Scale bars 4 µm, **(C)** Quantification of validated perinuclear ARC clusters per cell. ARC^S113–119A^ formed significantly fewer perinuclear clusters than ARC^WT^ (Mann–Whitney test, *p* = 0.0068). **(D)** Quantification of shuttling activity, expressed as the density of shuttling trajectories per total tracked molecules within each cell. Disruption of higher-order oligomerization significantly reduced nucleocytoplasmic shuttling (Mann–Whitney test, *p* = 0.0162). **(E)** Fraction of shuttling trajectories associated with validated perinuclear clusters. Approximately 32% of ARC^WT^ shuttling trajectories contacted a perinuclear cluster, whereas no cluster-associated shuttling events were detected for ARC^S113–119A^. **(F)** Representative ARC^WT^ perinuclear cluster (left) and example time series showing an individual ARC molecule repeatedly shuttling across the nuclear envelope while interacting with the cluster. Cyan arrows indicate nuclear entry and exit. Green lines show the trajectory. **(G)** Representative ARC^S113–119A^ cluster (left) and example time series illustrating the absence of cluster-associated nucleocytoplasmic shuttling.

To independently assess that the s113–119A mutation impairs higher-order oligomerization in clusters, we performed complementary selective time-resolved anisotropy with reversibly switchable states (STARSS), a fluorescence anisotropy approach that measures the rotational diffusion of protein assemblies and thereby reports their oligomeric state (35). Whereas SPT revealed the formation of perinuclear ARC clusters and their association with nucleocytoplasmic shuttling, STARSS analysis demonstrated significantly lower fluorescence anisotropy for ARC^s113–119A^ compared with ARC^WT^ specifically within ARC clusters, whereas no significant differences were observed in the nucleus or diffuse cytoplasm (Supplementary Fig. 3). These findings indicate that the s113–119A mutation selectively impairs higher-order (35)oligomerization within clustered ARC assemblies.

We next asked whether impaired cluster formation affected nucleocytoplasmic shuttling. Single-particle tracking revealed that neurons expressing ARC^s113–119A^ exhibited significantly fewer shuttling trajectories than neurons expressing than ARC^WT^ (Fig. 5D), indicating that ARC oligomerization promotes repeated nucleocytoplasmic shuttling. To further examine the relationship between clustering and shuttling, we quantified the proportion of shuttling trajectories associated with perinuclear clusters. In ARC^WT^-expressing cells, 32% of shuttling trajectories were associated with a perinuclear cluster, whereas no cluster-associated shuttling events were detected in cells expressing ARC^S113–119A^ (Fig. 5E).

Representative time-lapse sequences illustrate shuttling of individual ARC^WT^ particles associated with a perinuclear cluster, including both nuclear entry and subsequent nuclear exit (Fig. 5F). In contrast, the few puncta observed in ARC^s113–119A^ -expressing neurons remained largely stationary and were not associated with detectable nuclear shuttling events (Fig. 5G). Together, these findings demonstrate that ARC oligomerization promotes both perinuclear cluster formation and nucleocytoplasmic shuttling and identify perinuclear clusters as sites associated with shuttling ARC particles.

## Discussion

In this study, we provide the first direct evidence that ARC is a bona fide nucleocytoplasmic shuttling protein. By visualizing individual ARC particles in living neurons, we demonstrate that neuronal activity promotes dynamic bidirectional transport across the nuclear envelope. Furthermore, we identify activity-dependent perinuclear ARC clusters that are associated with shuttling trajectories and show that ARC oligomerization is required for both perinuclear cluster formation and nucleocytoplasmic shuttling. Together, these findings reveal a previously unrecognized mechanism linking ARC self-assembly to nucleocytoplasmic transport.

Previous studies established that ARC contains both nuclear localization and export sequences(15) and demonstrated that neuronal activity alters its nuclear localization(14, 15, 17). However, these approaches relied primarily on static measurements of protein distribution and therefore could not determine the trafficking behavior of individual ARC particles on per cell basis. By tracking ARC particles in living neurons, we demonstrate that ARC crosses the nuclear envelope in real-time. ARC crossings were low during unstimulated conditions and increased following neuronal stimulation (Fig. 1D-F). Consistent with the use of a constitutive promoter, we observed no change in the total number of ARC tracks following neuronal stimulation (Fig. 1G). By applying cLTP in neurons already expressing abundant ARC, our findings indicate that synaptic activity regulates the trafficking behavior of ARC independently of changes in its overall abundance.

This dynamic trafficking may have important implications for both the nuclear and synaptic functions of ARC. Das et al. recently proposed that newly synthesized ARC participates in an autoregulatory feedback loop by promoting a second wave of *Arc* transcription, thereby sustaining dendritic ARC mRNA and protein levels during prolonged synaptic plasticity (36). Although this model implicitly requires communication between the cytoplasmic and nuclear pools of ARC, the mechanism enabling this exchange has remained unresolved. By directly visualizing repeated nucleocytoplasmic shuttling of individual ARC particles, our findings identify a plausible transport mechanism through which cytoplasmic ARC could repeatedly access the nucleus, providing dynamic feedback in Arc transcription, chromatin remodeling, and nuclear speckle organization (14–19). More broadly, these findings suggest that nucleocytoplasmic trafficking may not only regulate the nuclear functions of ARC but also indirectly sustain its synaptic functions by coupling nuclear transcription with continued dendritic mRNA delivery and local translation during the later phases of synaptic plasticity.

Although considerable attention has been devoted to the spatial organization of ARC in dendrites (32), demonstrating that ARC assembles into diverse nanoscale structures at excitatory synapses(34) and accumulates in activity-dependent dendritic protein hubs maintained through repeated cycles of local translation (36), comparatively little is known about how neuronal activity reorganizes ARC within the soma. Our findings address this gap by demonstrating that neuronal stimulation redistributes ARC toward the perinuclear region, where it assembles into discrete clusters, identifying the perinuclear region as a previously underappreciated site of activity-dependent ARC organization.

Interestingly, the relationship between nucleocytoplasmic shuttling and perinuclear clustering was selective rather than universal. Specifically, only a subset of perinuclear clusters were associated with shuttling trajectories, whereas many shuttling trajectories did not contact detectable clusters (Figure 5E). Importantly, cluster–shuttling associations occurred more frequently than expected by chance (Figure 4D), indicating that these transport-associated clusters likely represent biologically meaningful sites rather than random spatial overlap. This interpretation is consistent with the emerging view of biomolecular condensates as dynamic and compositionally heterogeneous assemblies rather than uniform structures (37). Condensates sharing similar morphological features can differ substantially in molecular composition, RNA content, and protein interaction partners, enabling distinct biological functions despite their similar appearance. Accordingly, our findings raise the possibility that perinuclear ARC clusters comprise functionally specialized subpopulations.

Our findings further implicate ARC self-assembly in perinuclear organization and nucleocytoplasmic trafficking. ARC oligomerization encompasses a range of assembly states, with the dimer proposed to serve as a fundamental building block for the formation of higher-order oligomers. Purified ARC can assemble into higher-order species ranging from tetramers to larger assemblies, including particles containing approximately 32 ARC subunits and virus-like capsids comprising an estimated ∼120 subunits (6, 27, 28, 32, 34). Thus, higher-order oligomerization in this context does not refer exclusively to large capsid-like structures, but encompasses assemblies beyond the ARC dimer, including tetramers. Importantly, previous biochemical studies have shown that mutation of the N-terminal oligomerization motif spanning residues 113–119 impairs formation of these higher-order assemblies while preserving ARC dimers, an effect further supported by in situ nanobody-based proximity ligation analysis (28).

Although we did not directly determine the oligomeric state of ARC in the perinuclear clusters or shuttling trajectories, perturbation of this motif using ARC^s113–119A^ markedly reduced both perinuclear cluster formation and nucleocytoplasmic shuttling (Fig. 5C–D). This was supported by fluorescence anisotropy measurements showing altered self-association of ARC^s113–119A^ relative to ARC^WT^. Together with the previously established effect of this mutation on ARC oligomeric state, these convergent observations support a role for higher-order ARC assembly in perinuclear organization and nucleocytoplasmic trafficking. Perinuclear cluster formation was not completely abolished in ARC^s113–119A^-expressing neurons, indicating that higher-order assembly is unlikely to be the sole determinant of cluster formation. Notably, however, the remaining mutant clusters were not associated with detectable shuttling trajectories (Fig. 5E). This raises the possibility that higher-order ARC assembly is particularly important for the subset of perinuclear clusters associated with nucleocytoplasmic transport. Further work will be required to determine the oligomeric state and molecular composition of these transport-associated clusters and to establish whether higher-order assembly directly facilitates ARC trafficking across the nuclear envelope.

In sum, these findings suggest that the cytoplasmic and nuclear pools of ARC should not be viewed as independent populations but rather as dynamically interconnected through continuous nucleocytoplasmic exchange. Such dynamic trafficking, regulated in part by ARC self-assembly, provides a potential mechanism by which ARC can coordinate its diverse functions at synapses and within the nucleus following neuronal activity. Future studies should determine the molecular composition of these transport-associated perinuclear clusters and identify the mechanisms that distinguish them from non-transport-associated clusters.

### Limitations

Individual ARC trajectories were necessarily short owing to fluorophore photobleaching and the limitations of two-dimensional imaging, in which particles frequently moved out of the focal plane. Consequently, each trajectory represents a snapshot of ARC behavior rather than its complete transport history, and the frequency and duration of nucleocytoplasmic shuttling are likely underestimated. Finally, although stable expression of ARC from a constitutive promoter enabled us to isolate activity-dependent changes in transport and shuttling behavior, the trafficking behavior of endogenous ARC remains to be established.

## Methods

### Animal experiments

All experimental procedures approved by Norwegian National Research Ethics Committee in compliance with EU Directive 2010/63/EU, ARRIVE guidelines. Experiments were conducted by Federation of Laboratory and Animal Science Associations (FELASA) C course trained and certified researchers.

### Plasmids

For single-particle tracking, ARC^WT^-HaloTag and ARC^WT^-SNAP-tag constructs encoded full-length rat ARC fused at its C-terminus to HaloTag or SNAP-tag, respectively. Expression of both constructs was driven by the CMV promoter. The oligomerization-deficient ARC^s113–119A^-HaloTag construct was generated in the same CMV-driven backbone and contains alanine substitutions within the N-terminal oligomerization motif spanning residues 113–119. Disruption of this motif inhibits the assembly of ARC dimers into higher-order oligomers while preserving dimer formation (28).

For STARSS experiments, ARC had an AAV construct encoding either WT or s113–119A mutant with rsEGFP2 inserted between the N-terminal domain (NTD) and C-terminal domain (CTD) was used. Expression was driven by the EF-1α promoter, and the expression cassette was flanked by AAV2 inverted terminal repeats.

### Cell line

Human embryonic kidney 293FT (HEK293FT) cells, obtained from Thermo Fisher Scientific, were cultured in high-glucose Dulbecco’s Modified Eagle Medium (DMEM) (Sigma-Aldrich) and supplemented with 10% FBS. Upon reaching confluence, the cells were seeded onto 12 mm coverslips and transfected as described below.

### Primary neuronal cultures

The neuronal primary culture preparation was based on the protocols published for neuronal culture preparation and cryopreservation by Ishizuka et al. (38). Pregnant Wistar rats were deeply anesthetized and euthanized using CO₂ exposure. Embryos at embryonic day 18 (E18) were removed from the uterus and microdissected to isolate the hippocampus. The dissection was performed under a laminar airflow hood in cold Hank’s Balanced Salt Solution (HBSS) supplemented with 10 mM HEPES and 100 U/mL penicillin–streptomycin (Thermo Fisher Scientific). Following isolation, the hippocampi were maintained in cold HEPES buffer before undergoing tissue and cellular dissociation at 37 °C. Dissociation was performed using 0.05 % Trypsin–EDTA solution (Thermo Fisher Scientific) containing 10 mM HEPES and 100 U/mL penicillin–streptomycin. After enzymatic digestion, the tissue was mechanically dissociated by repetitive pipetting with a flame-polished glass Pasteur pipette. Finally, the dissociated cells were counted using a Neubauer chamber.

Glass coverslips (12 mm, Marienfeld-Superior, Lauda-Königshofen, Germany) were pre-treated with 1 N nitric acid (HNO₃) (Sigma-Aldrich, St. Louis, MO), followed by thorough washing with Milli-Q® sterile water. Neurons were then plated at a density of 4.0 × 10⁴ cells/cm² on coverslips coated with 1 mg/mL poly-D-lysine (Sigma-Aldrich). The neruons were initially cultured in Minimum Essential Medium (MEM) (Thermo Fisher Scientific) supplemented with 10% fetal bovine serum (FBS) (Sigma-Aldrich), 0.6% glucose (Sigma-Aldrich), 1 mM sodium pyruvate (Thermo Fisher Scientific), and 100 U/mL penicillin–streptomycin. Once the neurons adhered, the plating medium was replaced with Neurobasal™ medium (Thermo Fisher Scientific) supplemented with 2% B27™ (Thermo Fisher Scientific) and 0.25% GlutaMAX™-I (Thermo Fisher Scientific). Neurons were used for live-cell imaging between DIV 14–18 or fixed between DIV 18–21 using 4 % paraformaldehyde and 4 % sucrose in 0.1 M phosphate buffer (pH 7.5) for 20 minutes at room temperature. Fixed cells were processed for immunocytochemistry or proximity ligation assay.

### Chemical long-term potentiation (cLTP)

Chemically induced long-term potentiation of synaptic transmission (cLTP) was induced as described (25). Hippocampal neurons were incubated in Artificial Cerebrospinal Fluid (ACSF) without Mg^2+^ for 10 min. Then neurons were incubated in ACSF without Mg^2+^ supplemented with 20 µm Bicuculine, 3 μm Strychnine, and 200 μm Glycine for 10 min. The cells recovered in NBM+ for 2 h.

### Transfection and cellular labeling

Primary hippocampal neurons and HEK293FT cells were transfected with plasmids encoding ARC fused at the C-terminus to a HALO tag (ARC WT–HALO or ARC s113–119A–HALO) or with an empty-HALO control under the control of a CMV promoter. For dual-color SPT experiments performed at Janelia Research Campus, primary hippocampal neurons were co-transfected with ARC WT–SNAP and ARC s113–119A–HALO, allowing both ARC variants to be tracked within the same cell. ARC constructs contained the native 5′ and 3′ untranslated regions (UTRs). Transfections were performed using Lipofectamine 2000 (Invitrogen, L3000015). DNA and Lipofectamine were combined and incubated for 30 min at room temperature to allow complex formation. After 1 h of transfection, neurons were returned to their original conditioned Neurobasal medium supplemented with 0.25% GlutaMAX-I and maintained at 37°C in a humidified incubator with 5% CO₂. Twenty-four hours after transfection, neurons and HEK293FT cells expressing HALO-tagged constructs were labeled with 10 nM JF549 HaloTag ligand together with Hoechst or NucSpot nuclear dye for 30 min at 37°C (39). For dual-color SPT experiments, neurons co-expressing ARC WT–SNAP and ARC S113–119A–HALO were labeled with 6 nM JF646 SNAP-tag ligand and 5 nM JF549 HaloTag ligand for 30 min at 37°C. The labeling conditions produced sparse, spatially isolated fluorescent ARC particles that could be unambiguously localized and linked into trajectories during single-particle tracking. Following labeling, cells were washed four times for 10 min each with Neurobasal medium (neurons) or DMEM (HEK293FT cells) to remove unbound ligand.

HEK293FT cells were fixed for downstream analysis. Neurons were imaged either under unstimulated conditions or following chemical long-term potentiation (cLTP) stimulation. For cLTP experiments, neurons were labeled 50 min after stimulation. Imaging was performed in Mg²⁺-free artificial cerebrospinal fluid (ACSF).

### Image setup for single-particle tracking

Single-particle tracking (SPT) experiments were performed at two imaging sites using Nikon Ti2-based inverted microscope platforms. The two systems differed in effective pixel size (0.11 or 0.16 µm pixel⁻¹) and field of view (256 × 256 or 216 × 216 pixels). Movies were acquired for up to 5,000 consecutive frames at a 50 ms exposure time (20 Hz). For recordings containing fewer frames, analyses were performed using the full available movie. Recording duration was determined individually for each movie and incorporated into downstream analyses where appropriate. All spatial measurements were converted using microscope-specific pixel calibration during image analysis.

University of Bergen: Live hippocampal neurons were imaged using a Nikon Eclipse Ti2 inverted microscope equipped with a 60× oil-immersion CFI Apo TIRF objective (NA 1.49, WD 0.12 mm) and a Prime BSI Express scientific CMOS camera (Teledyne Photometrics). Single-particle imaging was performed using highly inclined and laminated optical sheet (HILO) illumination. Excitation was provided by 561 nm (60 mW) or 640 nm (50 mW) laser lines operated at full laser power, depending on the fluorophore used. Images were acquired at 16-bit depth with 1×1 binning using a 50 ms exposure time (20 Hz acquisition rate). Image sequences were recorded as 256 × 256 pixel regions of interest to maximize temporal resolution. Live-cell imaging was performed at 37°C using an OkoLab stage-top incubation system with CO₂ control and objective heating. The Nikon Perfect Focus System (PFS) was used to maintain focal stability throughout time-lapse acquisition.

Janelia Research Campus: Live hippocampal neurons were imaged using a Nikon Ti2 inverted microscope configured for objective-type TIRF illumination. Images were acquired using a 100× oil-immersion objective (NA 1.49) and an Andor DU-897 EMCCD camera (Andor Triple DU-897 system) at 16-bit depth with 1×1 binning. Time series were recorded as 216 × 216 pixel regions of interest for 5,000 consecutive frames with a 50 ms exposure time (20 Hz acquisition rate). Dual-color imaging was performed exclusively at Janelia using sequential excitation with 561 nm and 647 nm laser lines operated at full laser power. The EM gain was set to 300 for both channels. TIRF illumination was achieved using an incidence angle of 49°, and the Nikon Perfect Focus System (PFS) was used to maintain focal stability throughout time-lapse acquisition.

### Image analysis

#### Single-particle detection and trajectory reconstruction

Single-particle tracking (SPT) was performed in MATLAB using TrackIt (40) together with custom MATLAB scripts (see code availability). aw image sequences were processed using a four-step analysis pipeline. First, single-particle signals were enhanced using two wavelet filters (41). Candidate particles were then identified by local-maxima detection with an intensity threshold of 2, localized by two-dimensional Gaussian fitting using the TrackNTrace fast-fitting algorithm, and linked into trajectories using a nearest-neighbour tracking algorithm. Tracking parameters were set manually and kept constant across all conditions: tracking radius = 0.8672 µm, a minimum trajectory length of five frames, a maximum gap length of five frames, and a minimum trajectory length of five frames before gap closing was permitted. To avoid transient over-brightness immediately following image acquisition, the first 50 frames of each movie were excluded from analysis.

### Nuclear border definition and crossing analysis

The nucleus was visualized using Hoechst or NucSpot 650 staining. For each movie, the nuclear boundary was manually delineated from the nuclear fluorescence image before trajectory analysis to generate a binary nuclear mask.

To minimize false-positive transport events arising from localization uncertainty at the nuclear envelope, nucleocytoplasmic transport was analysed using a custom MATLAB pipeline implementing a strict core-compartment approach. Two non-overlapping core compartments were generated from the nuclear mask: a nuclear core comprising pixels located at least 0.2 µm inside the nuclear boundary and a cytoplasmic core comprising pixels located at least 0.2 µm outside the nuclear boundary. Localizations within the ±0.2 µm border region were not assigned to either compartment and were excluded from transport classification. Microscope-specific pixel calibration was applied before evaluating each localization relative to the nuclear mask.

Trajectories containing fewer than five localizations or fewer than two retained core localizations were excluded from transport analysis. Each retained localization was assigned to either the nuclear or cytoplasmic core according to its spatial position. A strict nuclear crossing event was defined as a transition between the cytoplasmic and nuclear core compartments in consecutive retained localizations (cytoplasm → nucleus or nucleus → cytoplasm). The total number of compartment transitions was determined for each trajectory.

Trajectories exhibiting a single compartment transition were classified as nuclear import or export according to the direction of the transition. Trajectories exhibiting two or more compartment transitions were classified as shuttling. For shuttling trajectories, the first and last retained core localizations were used to classify the trajectory as nucleus-to-nucleus, nucleus-to-cytoplasm, cytoplasm-to-nucleus, or cytoplasm-to-cytoplasm. All analysis parameters were fixed before analysis and applied identically to all cells, datasets, and experimental conditions.

Track density was calculated as the number of analysed trajectories normalized to movie duration and imaging area and is reported as trajectories min⁻¹ µm⁻².

### Mean intensity analysis

To quantify the subcellular distribution of ARC, custom MATLAB scripts were used to measure fluorescence intensity within manually defined nuclear, perinuclear, and cytoplasmic compartments. Nuclear and cytoplasm boundaries were manually delineated from the nucleus reference image (Hoecht staining) and cell outline using average intensity, respectively. A perinuclear compartment was generated by expanding the nuclear mask by 2.0 µm and subtracting the nuclear region. To ensure that only intracellular signal was included, the perinuclear ring was clipped to the manually defined cell ROI, and the remaining cytoplasmic compartment was defined as the cell ROI excluding both the nucleus and the clipped perinuclear ring.

For each image frame, background fluorescence was estimated as the fifth percentile of pixel intensities across the image and subtracted prior to analysis. Pixels approaching detector saturation were excluded. Mean or median fluorescence intensity was calculated within each compartment, with median intensity used for all analyses presented in this study. Mean values across all image frames were calculated for each cell and used for statistical analysis. Quality-control images showing the generated compartment masks were produced for each analyzed cell.

### Identification of perinuclear ARC clusters

Perinuclear ARC clusters were identified using a custom MATLAB pipeline implementing density-based spatial clustering of applications with noise (DBSCAN). Tracked single-particle localizations within the perinuclear compartment were pooled for each cell. The perinuclear compartment was defined as the region extending from 0.2 to 2.0 µm outside the nuclear boundary and clipped to the manually delineated cytoplasmic ROI to exclude extracellular regions. All spatial measurements were converted using microscope-specific pixel calibration. Localizations were clustered using DBSCAN with a search radius (ε) of 0.25 µm and a minimum cluster size of 20 localizations. Adjacent DBSCAN fragments separated by less than 0.25 µm were merged to reconstruct contiguous clusters. Candidate clusters were filtered using predefined spatial and intensity criteria. Clusters were required to have an area between 0.01 and 5.0 µm² and to be supported by localizations originating from at least three independent trajectories. Fluorescence intensity was evaluated on the average intensity image following background subtraction using the fifth percentile image intensity. Candidate clusters were accepted only if the 90th percentile cluster intensity exceeded a four-fold enrichment relative to the local perinuclear background and exceeded the local background by at least 2.5 median absolute deviations (MAD)

All clustering parameters were empirically optimized during method development and subsequently fixed before analysis. The same parameter values were applied to all datasets, cells, and experimental conditions without modification to ensure unbiased comparisons.

### Association between shuttling trajectories and validated clusters

Only trajectories classified as shuttling by the strict compartment-transition analysis were included in cluster association analyses. Validated cluster masks were expanded by 0.35 µm to account for localization uncertainty. A shuttling trajectory was classified as cluster-associated when at least one localization intersected the expanded validated cluster mask. Direct overlap with the original cluster mask was also recorded. For each cell, the total number of shuttling trajectories, the number contacting validated clusters, and the percentage of cluster-associated shuttling trajectories were calculated.

### Correlation analysis

To determine whether cluster abundance was associated with cluster-dependent nucleocytoplasmic transport, the relationship between the number of validated clusters per cell and the fraction of shuttling trajectories contacting validated clusters was assessed using Spearman rank correlation. Cells without detected shuttling trajectories were excluded because the fraction of cluster-associated shuttling could not be defined. Data from the Norway and Janelia datasets were pooled after confirming identical analysis parameters, and each cell contributed a single observation to the correlation analysis.

### Cluster randomization analysis

To determine whether nucleocytoplasmic shuttling occurred more frequently at validated ARC clusters than expected by chance, observed cluster-associated shuttling events were compared with randomized expectations generated by a custom MATLAB analysis pipeline. For each validated cluster, the observed number of shuttling trajectories contacting the cluster was compared with the expected number obtained from repeated randomization while preserving the overall properties of the experimental trajectories. The randomized distribution was used to calculate the expected mean number of contacts and an empirical one-sided probability for observing an equal or greater number of contacts by chance. Clusters were classified as active when contacted by at least one observed shuttling trajectory. Active clusters were further classified as significantly above chance when the observed number of contacting trajectories exceeded the randomized expectation and the empirical upper-tail probability was less than 0.05. Observed and randomized contact frequencies were compared using paired Wilcoxon signed-rank tests performed separately for all validated clusters and for active clusters only. Cluster-level analyses report both the number of clusters and the number of contributing cells.

### Quantification and statistical analysis

Statistical analyses were performed using GraphPad Prism (version 10.6.1). Data are presented as mean ± s.e.m. unless otherwise stated. Statistical tests used for individual experiments are indicated in the corresponding figure legends and included Mann–Whitney U tests, paired Wilcoxon signed-rank tests, Friedman tests with Dunn’s multiple-comparisons correction, Spearman rank correlation, and unpaired t-tests where appropriate. Statistical significance was defined as p < 0.05.

## Acknowledgements

Thanks to Dr. Deepika Walpita for preparing hippocampal neurons at Janelia Research Campus, and Dr. Giovanna Coceano for amplifying and sequencing plasmids at Karolinska Institute.

## Supplementary

**Supplementary Figure 1. Validation of JF549 labeling of ARC-HALO constructs.**

**(A)** Representative fluorescence images of cells expressing empty-HALO, ARC^WT^-HALO, or ARC^s113–119A^-HALO following immunostaining for ARC (green) and labeling with the JF549 HaloTag ligand (red). Merged images show spatial overlap between ARC immunoreactivity and JF549 fluorescence for both ARC^WT^-HALO and ARC^S113–119A^-HALO, whereas empty-HALO-expressing cells show JF549 fluorescence in the absence of detectable ARC immunoreactivity.

**(B-C)** Pixel-wise correlation between ARC antibody fluorescence and JF549 fluorescence for ARC^S113–119A^-HALO (B) and ARC^WT^-HALO (C). Pearson correlation coefficients were calculated for individual cells, with an average Pearson correlation coefficient of 0.81 for ARC^S113–119A-HALO (n = 9 cells) and 0.79 for ARC^WT-HALO (n = 8 cells).

**Supplementary figure 2. Empty-HALO controls for ARC nuclear crossing and perinuclear cluster detection.**

**(A–B)** Representative examples of trajectories detected in neurons expressing empty-HALO under unstimulated conditions (A) and 2 h following cLTP (B). Average-intensity projections and the nuclear border are shown, with detected nuclear crossing trajectories indicated in different colors.

**(C)** Fraction of empty-HALO trajectories undergoing strict nuclear crossing events per cell under unstimulated conditions and 2 h following cLTP. No significant difference was observed between conditions (Mann–Whitney test).

**(D)** Empty-HALO trajectory density per cell (tracks min⁻¹ µm⁻²) under unstimulated conditions and 2 h following cLTP. No significant difference was observed between conditions (Mann–Whitney test). Together, these data indicate that the activity-dependent increase in nuclear crossing observed for ARC-HALO is not reproduced by empty-HALO.

**(E–F)** Representative images of neurons expressing empty-HALO (E) or ARC-HALO (F) under unstimulated conditions, illustrating the fluorescence images used for perinuclear cluster detection.

**(G)** Number of validated perinuclear clusters per cell in unstimulated neurons expressing empty-HALO or ARC-HALO. No significant difference was observed between groups (Mann–Whitney test).

**(H–I)** Representative images of neurons expressing empty-HALO (H) or ARC-HALO (I) 2 h following cLTP. ARC-HALO displayed prominent perinuclear clusters following stimulation, whereas these structures were largely absent in empty-HALO-expressing neurons.

**(J)** Number of validated perinuclear clusters per cell 2 h following cLTP in neurons expressing empty-HALO or ARC-HALO. Significantly more clusters were detected in ARC-HALO-expressing neurons than in empty-HALO controls (Mann–Whitney test, **P < 0.01). These results support that the activity-dependent perinuclear clusters detected by the analysis pipeline are specific to ARC rather than arising from HaloTag labeling or nonspecific detection by the cluster-analysis pipeline.

**Supplementary Figure 3. STARSS analysis reveals reduced higher-order oligomerization of ARC^S113–119A^ within ARC clusters.**

**(A–B)** Representative fluorescence images of HeLa cells expressing ARC^WT (A) or the oligomerization-deficient ARC^S113–119A mutant (B) used for selective time-resolved anisotropy with reversibly switchable states (STARSS) analysis. Scale bars, 10 µm.

**(C)** Fluorescence anisotropy measured within ARC clusters was significantly lower for ARC^S113–119A^ compared with ARC^WT^ (****P < 0.0001, Mann-Whitney test), consistent with reduced higher-order oligomerization of the mutant within clustered ARC structures.

**(D)** Comparison of fluorescence anisotropy measured within the nucleus, ARC clusters, and cytoplasm. In ARC^WT^-expressing cells, anisotropy was significantly higher within ARC clusters than in either the nucleus or cytoplasm (****P < 0.0001, Friedman test with Dunn’s multiple-comparisons test), whereas no significant differences between compartments were detected for ARC^S113–119A^. Together, these results indicate that higher-order ARC organization is selectively associated with clustered structures and is impaired by the S113–119A mutation.

