## Supplementary figures and images for "Single-particle tracking reveals neuronal activity-dependent shuttling of ARC/ARG3.1 protein between cytoplasmic clusters and the nucleus"

### Supplemental data

Supplementary Figure 1

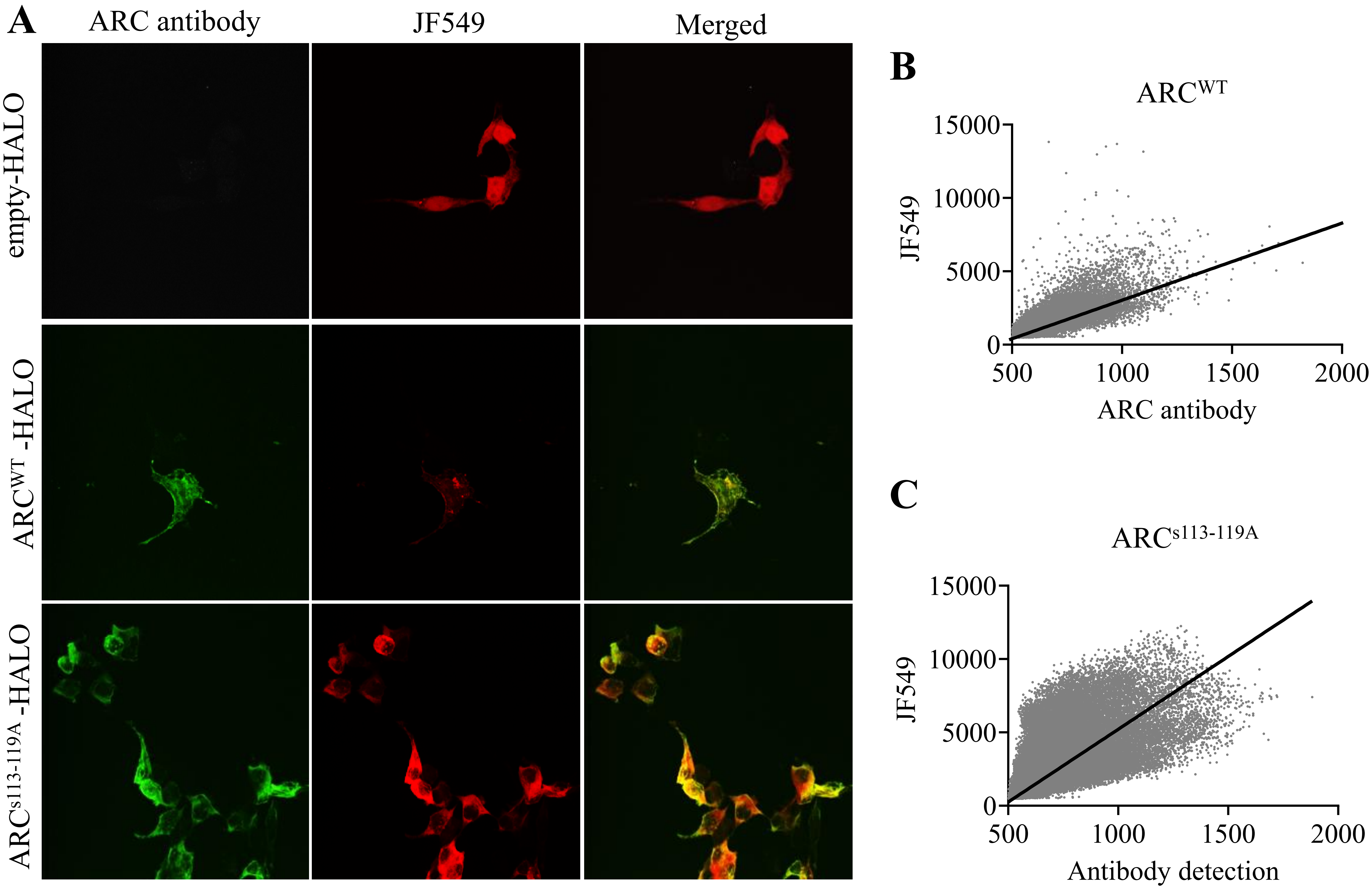

Supplementary Figure 2

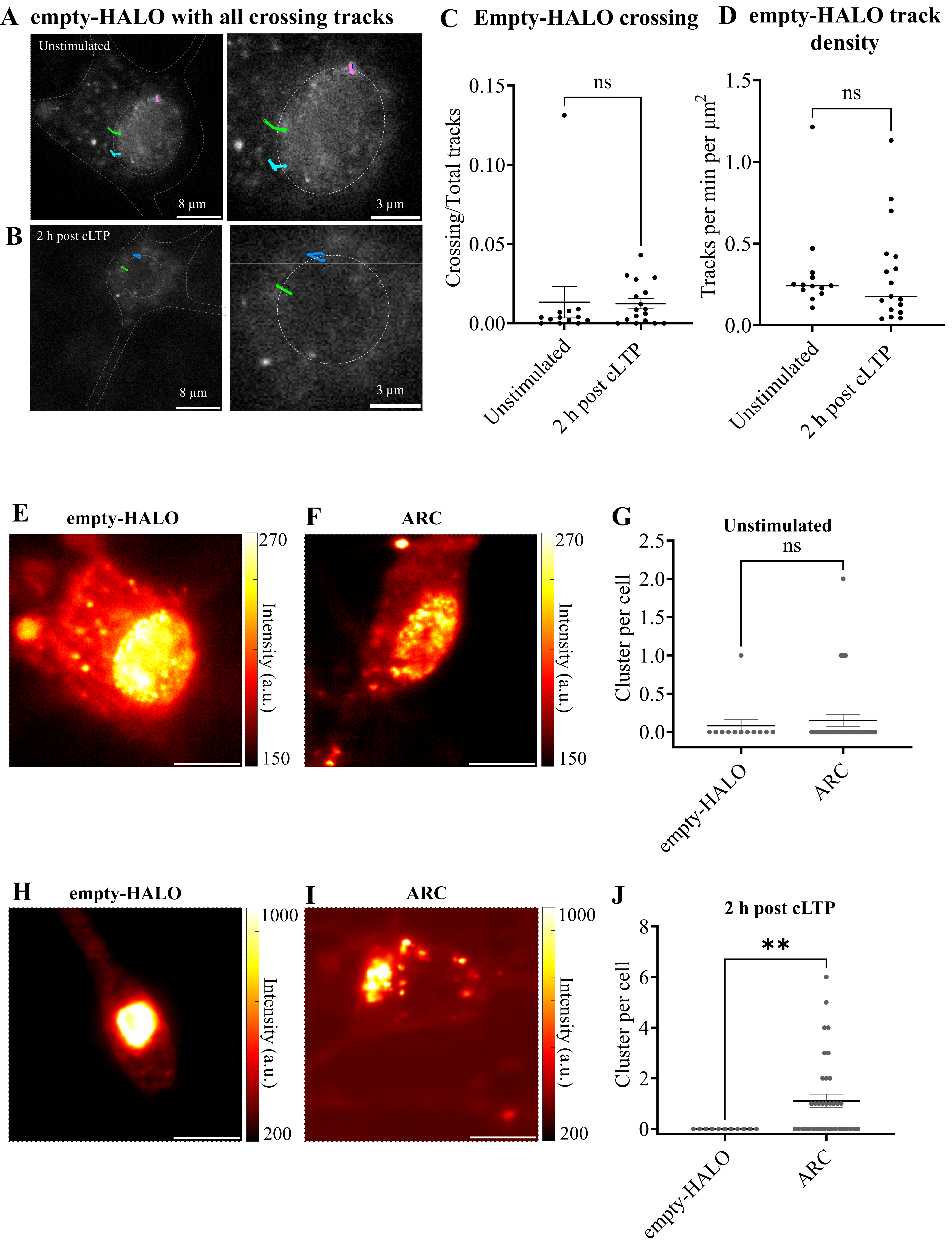

Supplementary Figure 3

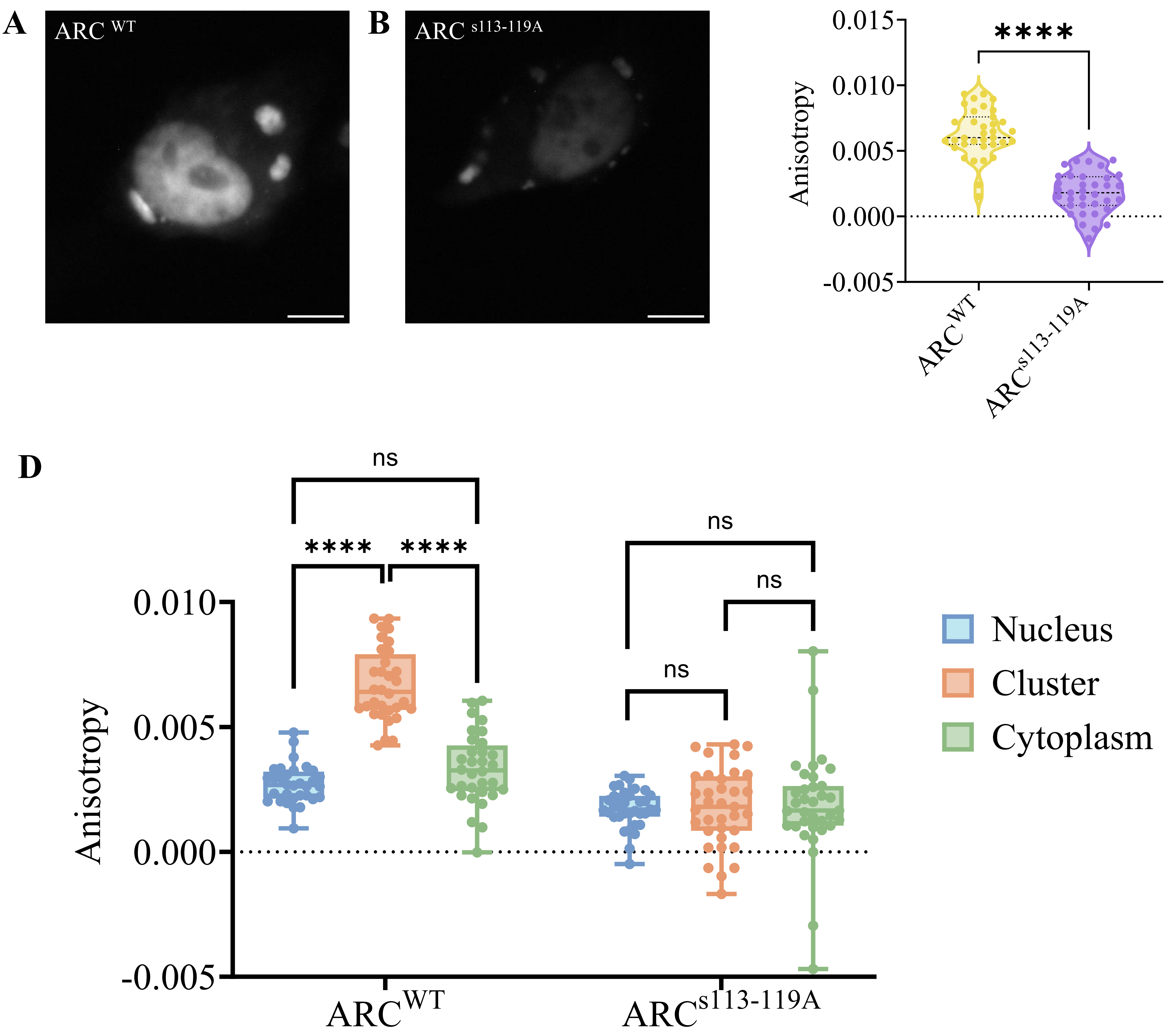
